# Dynamic bistable switches enhance robustness and accuracy of cell cycle transitions

**DOI:** 10.1101/2020.08.11.246017

**Authors:** Jan Rombouts, Lendert Gelens

**Affiliations:** Laboratory of Dynamics in Biological Systems, Department of Cellular and Molecular Medicine, University of Leuven (KU Leuven), B-3000 Leuven, Belgium

## Abstract

Bistability is a common mechanism to ensure robust and irreversible cell cycle transitions. Whenever biological parameters or external conditions change such that a threshold is crossed, the system abruptly switches between different cell cycle states. Experimental studies indicate that the shape of the bistable response curve changes dynamically in time. Here, we show how such a dynamically changing bistable switch can provide a cell with better control over the timing of cell cycle transitions. Moreover, cell cycle oscillations built on bistable switches are more robust when the bistability is modulated in time. Our results are not specific to cell cycle models and may apply to other bistable systems in which the bistable response curve is time-dependent.

**Author summary:** Many systems in nature show bistability, which means they can evolve to one of two stable steady states under exactly the same conditions. Which state they evolve to depends on where the system comes from. Such bistability underlies the switching behavior that is essential for cells to progress in the cell division cycle. A quick switch happens when the cell jumps from one steady state to another steady state. Typical of this switching behavior is its robustness and irreversibility. In this paper, we expand this viewpoint of the dynamics of the cell cycle by considering bistable switches which themselves are changing in time. This gives the cell an extra layer of control over transitions both in time and in space, and can make those transitions more robust. Such dynamically changing bistability can appear very naturally. We show this in a model of mitotic entry, in which we include a nuclear and cytoplasmic compartment. The activity of a crucial cell cycle protein follows a bistable switch in each compartment, but the shape of its response is changing in time as proteins are imported and exported from the nucleus.

## 1 Introduction

Multistability is one of the clearest manifestations of nature’s inherent nonlinearity. A multistable system can, under exactly the same conditions, be in different stable steady states. Consider a ball moving on a hilly terrain under the influence of gravity, where every valley corresponds to a stable state for the ball (Fig. 1A). When there are multiple valleys, the ball’s initial position determines where it will end up. These valleys can appear and disappear as the shape of the terrain changes. Another way of looking at such a changing terrain is obtained by plotting the steady state position of the ball (labeled output) in function of a parameter that determines the shape of the terrain (labeled input) (Fig. 1B). Here, for low input, there is only one steady state (situation 1). By increasing the parameter, a new state appears and the system is said to be bistable (situation 2). When the input crosses a threshold value, the initial stable valley disappears and the ball is forced to move to the right valley (situation 3). This transition is discontinuous, fast and irreversible. In between two stable states, there is an unstable steady state (the maximum in Fig. 1A and the dashed line in Fig. 1B). The points at which the stable and unstable steady state coalesce define the threshold values. These points are also called saddle-node points in the language of bifurcation theory.

**Fig 1.**
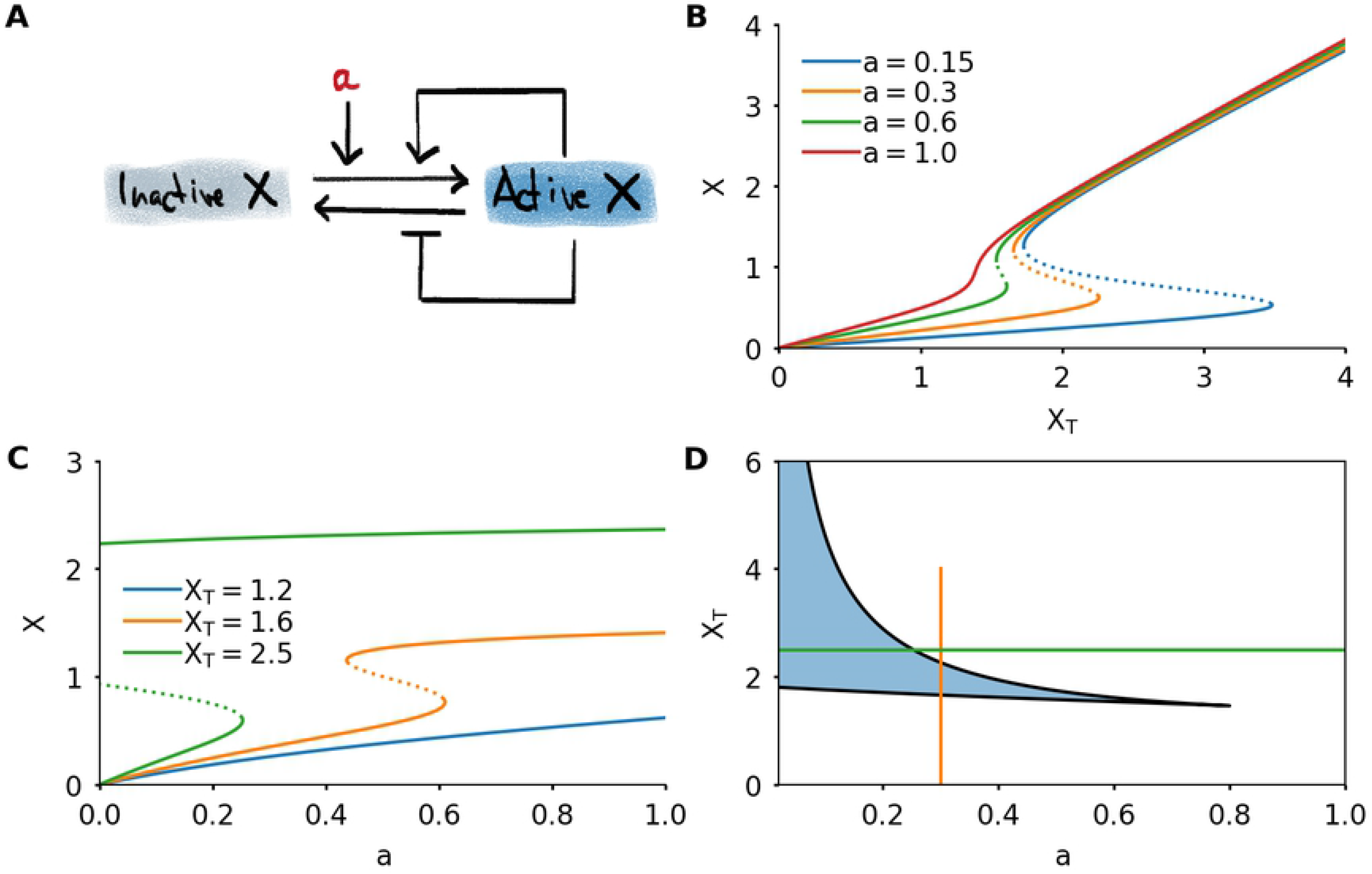
Bistability allows robust switching and is common in the cell cycle. A) A clear example of bistability in a dynamical system occurs when a little ball moves under the influence of gravity on a hilly terrain. Valleys correspond to stable steady states. These can be created and destroyed under influence of an external parameter. When a steady state disappears, the ball quickly transitions to another steady state. B) Representation of the ball’s position as function of a parameter which determines the shape of the terrain in Panel A. When the input increases beyond a threshold, the left equilibrium position in Panel A disappears and the ball quickly moves to the other stable position. C) In the cell cycle, bistable switches underlie some of the important transitions and checkpoints.

This simple mechanical example has equivalents in all sorts of physical and biological systems, where Newton’s laws of motion and the hilly terrain are replaced by chemical reactions, predator-prey interactions, heat transport or other mechanisms. In climate and ecology studies, transitions to a new steady state are often called tipping points [1, 2], and they are of special interest given current climate change. Bistability is present on all scales, ranging from the global climate system [2] to a single cell [3]. The genetic system involving the *lac* operon in an *E. coli* cell [4] allows bacteria to switch between using glucose or lactose. Bistability in actin polymerization state enables a cell to quickly and robustly switch between different migration modes [5]. Besides these *in vivo* examples, bistable responses have also been observed in purified kinase-phosphatase systems [6, 7] and are a common objective in the design of synthetic genetic systems [8]. The concept of bistability and irreversible transitions also plays an important role in cell differentiation. There, the image of balls rolling down valleys is echoed in Waddington’s epigenetic landscape [9].

On the molecular level, bistability is generated by the interplay of a large amount of molecules participating in chemical reactions. The conditions under which these reaction networks generate bistability have been extensively studied. Typically, one needs highly nonlinear (ultrasensitive) responses and positive feedback loops [10, 11]. However, bistability can also be present in simple systems with a minimal amount of components governed by mass-action kinetics. Finding the conditions under which such systems generate multistability is one of the important questions asked in chemical reaction network theory, where mostly algebraic methods are used to analyze these systems (eg. [12–15]).

To survive and proliferate, a cell has to replicate its DNA and structural components, and then distribute this material evenly to its daughters. This process is governed by the orderly progression through different phases of the cell cycle. The eukaryotic cell cycle contains various checkpoints and transitions in which bistability plays a role (Fig. 1C), and can even be viewed as a chain of sequentially activated bistable switches [16–18]. These switches provide robustness and directionality to the cell cycle and ensure the genome’s integrity. Both the ‘commitment point’, where a cell becomes committed to enter the cell cycle, and the transition from G1 to S phase have been associated to underlying bistable switches [19–22]. Later, after the cell has duplicated its DNA, there is a sudden transition from G2 to mitosis, characterized by the prompt activation of cyclin-dependent kinase 1 (Cdk1). This sudden mitotic entry has also been shown to be controlled by two bistable switches [23–26]. Further in mitosis, the spindle assembly checkpoint (SAC) controls the correct separation of sister chromatids at the metaphase-anaphase transition [27]. Theoretical models have shown that there are molecular mechanisms that can lead to bistability underlying this checkpoint [28–30].

The standard view of all these cell cycle transitions is that the two states are represented by two branches of a static bistable response curve. The transition happens when a slowly changing input reaches a threshold, upon which the system jumps to the other branch of the curve (Fig. 1B). Throughout this work we will focus on the bistable switch in mitotic entry that has been arguably best characterized experimentally and theoretically. Already in the early 1990s, mathematical models showed how biochemical interactions could lead to cell cycle oscillations that switched between interphase and mitosis [31, 32], and later predicted that bistability might be at the basis of the mitotic entry transition [33, 34]. This bistability was later verified experimentally [23, 24]. The kinase Cdk1 becomes active when bound to a Cyclin B subunit, and is involved in two feedback loops: Cdk1 activates the phosphatase Cdc25, which removes an inhibitory phosphorylation on Cdk1, thereby activating it and closing a double positive feedback loop. Secondly, Cdk1 inhibits Wee1, a kinase responsible for inhibiting Cdk1 through phosphorylation. This constitutes a double negative feedback loop. Due to ultrasensitivity in these feedbacks, a bistable response of Cdk1 activity to Cyclin B concentrations is generated. The shape of this switch depends, among others, on the amounts of Wee1 and Cdc25 present [35] (for more details, see Section 2.1).

In many cell cycle transitions, however, the parameters which determine the shape of the switch are also changing, either slowly or in a more sudden fashion. Typically the shape of the bistable response curve depends on the total concentration of proteins implicated in the feedback loops. These concentrations may change, either due to production and degradation, or due to relocalization of proteins in space. We can consider the cell as a set of compartments with slow fluxes between them. If each compartment is well-mixed, it has its own bistable response curve. The shape of this curve depends on the concentrations of proteins in that compartment, which can change over time as proteins relocalize. Note that compartmentalization has been studied in the context of bistability already: adding different compartments can be a mechanism of generating a bistable response, where there is none in a single well-mixed system [36, 37]. In Section 2.1, we discuss how considering nucleus and cytoplasm as compartments can alter the bistable switches governing mitotic entry. The importance of dynamically changing the bistable response curve has been acknowledged before, mostly in the context of lowering an activation threshold. For example, the threshold for mitogenic signaling, which defines the commitment point, can be influenced by DNA damage, cell volume or cell contacts [22, 38]. This provides extra control over the timing of passing the commitment point. At mitotic entry, Wee1 is known to be quickly degraded [39, 40], which lowers the threshold for Cdk1 activation [41] and triggers a transition into mitosis.

Bistability also lies at the heart of an important class of oscillations which appear time and again in chemical, biological and physical systems. These oscillations are called relaxation oscillations, and consist of slow progress along the branches of a bistable system, with sudden jumps between them. Eminent examples of relaxation oscillators are the Van der Pol and FitzHugh-Nagumo type systems. Whereas they were developed as models of electrical systems, either engineered, or in neurons, now they are often used as generic oscillating systems which can exhibit different kinds of dynamics [42]. Nonlinear oscillators generate many periodic phenomena in cell biology, among which circadian rhythms, metabolic oscillations, and also the embryonic cell cycle (see the books [43, 44] and review papers [45, 46] for overviews of biological oscillations). The embryonic cell cycle – in contrast to the somatic cycle – largely lacks checkpoint control, gap phases and even growth. The cycle is driven forward as a true oscillator by periodic production and degradation of proteins.

Here, we investigate how dynamically changing bistable switches affect transitions and relaxation oscillations. To motivate our studies, we first show how including different cellular compartments in a model of mitotic entry leads naturally to a situation with bistable switches that change in time. Next, we explore the concept of dynamically changing switches using a simple model. After introducing the model, we discuss how a single transition, such as the crossing of a cell cycle checkpoint, is affected by dynamically changing the activation threshold and the shape of the response curve. Moreover, we describe a mechanism which may be at play in spatially extended systems. There, bistability can lead to traveling fronts, whose speed depends on the shape of the bistable response curve. Front propagation can therefore dynamically change as proteins – which determine the shape of the response curve – are redistributed in space. We then discuss how such a dynamically changing switch affects relaxation oscillations. At each point we interpret our general results in the context of mitotic entry.

## 2 Results

### 2.1 A model for mitotic entry shows how dynamic switches appear in two cellular compartments

Mitotic entry is triggered by the activation of the kinase Cdk1, which sets into motion many of the changes a cell undergoes during mitosis. Cdk1 becomes active when bound to a Cyclin B subunit. Additionally, Cdk1 activity is controlled by its phosphorylation state, which is regulated by the kinase Wee1 and the phosphatase Cdc25 (Fig. 2A). In turn, Cdk1 itself activates Cdc25 and inactivates Wee1. These feedback loops produce a bistable response of Cdk1 activity as function of total Cyclin B levels [23, 24], very similar to the model we use in the rest of the paper. The different feedback loops have been characterized in detail [47, 48]. Mitotic entry involves many other mechanisms, such as a phosphatase switch [25] or the regulation of other kinases such as those from the Polo or Aurora families. An excellent recent review of the mitotic entry transition is given by Crncec and Hochegger [49].

**Fig 2.**
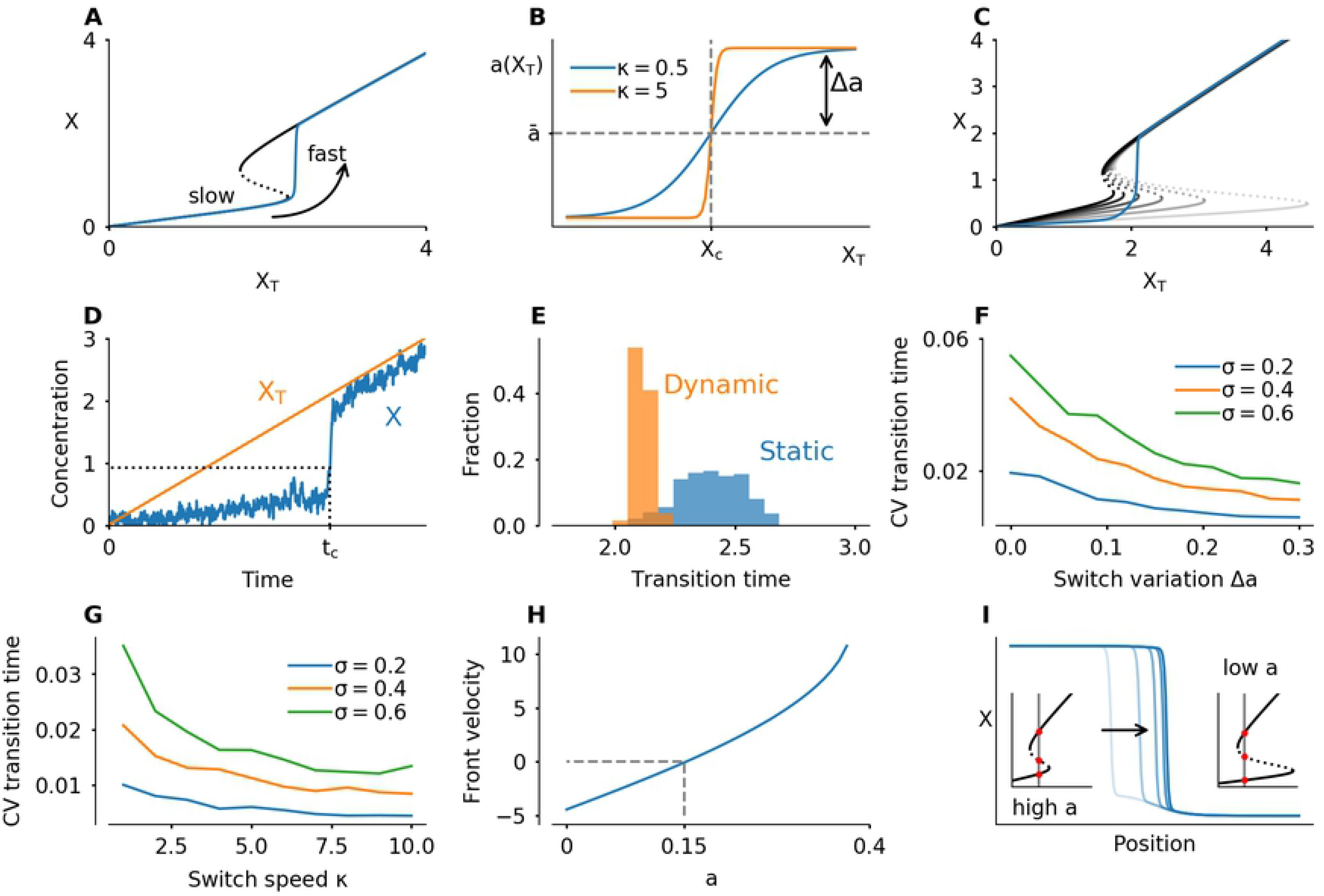
A model for mitotic entry shows how dynamic switches appear in two cellular compartments. A) The core protein interaction network involved in mitotic entry with double positive and double negative feedback loops centered on the Cyclin B-Cdk1 complex. Additional feedbacks act on the import rates of Cdc25 and Cyclin B-Cdk1. B) Two cellular compartments can exchange proteins through import and export. The total concentration of Cdc25 in each compartment determines the shape of the bistable response, which changes over time. C) Simulation of mitotic entry driven by Cyclin B production in the cytoplasm. Left: initially the activation threshold in the nucleus lies to the far right, due to the dominance of Wee1 over Cdc25 there. The activation threshold shifts left as Cdc25 is imported, which happens faster as Cdk1 activity rises in the cytoplasm. Middle: the activation threshold for Cdk1 activation is first crossed in the cytoplasm. The sudden jump in Cdk1 activation effects a sudden increase of Cdc25 import into the nucleus, which in turn quickly lowers the activation threshold there. Right: the decrease of the threshold in the nucleus triggers activation of Cdk1, leading to additional import of Cyclin B and a high Cdk1 activity. The black dot denotes the position of the system, the red curve corresponds to the bistable response at a given time point whereas the gray lines are snapshots of the bistable response at times leading up to this point. An animation which more clearly illustrates the dynamics can be found in S4 Video.

One particular source of additional regulation comes from the spatial localization of the different proteins. In mitosis the Cyclin B-Cdk1 complexes accumulate in the nucleus [36, 50]. Cdc25 also translocates to the nucleus at mitotic entry [51]. Wee1, the kinase inhibiting Cdk1, is mostly nuclear during interphase [52], possibly to make sure that Cdk1 is not activated too early, i.e. before DNA replication – which takes place in the nucleus – is complete. Spatial regulation of other mitotic regulators such as Polo [53] and Greatwall [54, 55] has recently been shown to be important for correct progress of mitosis as well.

All of these spatial translocations influence the behavior of the system, and here we show that this can be interpreted in the framework of a bistable switch with dynamically changing shape. To this end we extend the cell cycle model of Yang and Ferrell [56] to include two different compartments: the nucleus and the cytoplasm. In each compartment, Cdk1 activation is governed by the feedback loops through Wee1 and Cdc25. In addition, proteins can move into and out of the nucleus. The nuclear import rates may depend on the concentrations of Cdk1, to include spatial feedback [36]. In our simplified version, we assume that active cytoplasmic Cdk1 enhances import of Cdc25 and nuclear Cdk1 enhances import of Cyclin B. These assumptions are approximations of the experimentally known feedbacks [36, 51, 53]. We assume that Wee1 concentrations are higher in the nucleus. If the import and export rates are slow relative to the activation dynamics of Cdk1, we can consider each compartment to be nearly in steady state. This steady state, in turn, follows the bistable response curve of Cdk1 as a function of total Cyclin B. Due to translocation of Cdc25, the shape of these curves varies (Fig. 2B). We do not aim to include all of the complexity of mitotic entry described in the previous paragraph. Rather, we want to show how including a minimal spatial component using plausible mechanisms leads to changed mitotic entry dynamics, which can be interpreted using the dynamic bistable switches. More details and the full set of equations used can be found in the Methods section.

Adding these compartments leads to a mitotic entry in different steps (Fig. 2C). First, Cyclin B accumulates in the cytoplasm. At the start, the threshold for Cdk1 activation is lower in the cytoplasm than in the nucleus, due to the lower Wee1 concentration there. As a consequence, Cdk1 activation occurs first in the cytoplasm. This activation triggers the import of Cdc25 in the nucleus, which lowers the activation threshold there and allows Cdk1 activation in the nucleus. This triggers a translocation of even more Cyclin B to the nucleus. These two effects ensure that the activation of Cdk1 in the nucleus is very fast, irreversible, and happens after cytoplasmic activation of Cdk1. Once Cdk1 is activated in the nucleus, nuclear import of Cyclin B stays high. Most newly synthesized Cyclin B will be imported in the nucleus, further raising Cdk1 activity levels. Cdk1 activity in the cytoplasm settles at a nearly constant value. The animation S4 Video makes the time evolution of the different switches more clear.

The key observation we want to stress with this illustration is that the activation threshold is different in nucleus and cytoplasm, and importantly, that this threshold is controlled by translocation of Cdc25. By spatially regulating Cdc25, the cell has an additional layer of control over the timing of Cdk1 activation. The combined feedbacks lead to quick Cdk1 activation in the nucleus, and an enhanced import rate makes sure that Cdk1 activity in the nucleus increases further.

This example can be extended by including nuclear envelope breakdown (NEBD). Cdk1 activation triggers this event, which effectively mixes the two compartments. In turn, the two bistable response curves collapse to a single one. This provides another example of dynamically changing bistable switches. Moreover, the translocation of other proteins, such as the kinase Greatwall, will likely have a similar effect on the shape

### 2.2 A simple model produces bistability in protein activity

The results of the previous section show that a dynamically changing switch can appear naturally in a biochemical system. To investigate the consequences of such a changing switch in more depth, we introduce a simple model. The model describes a protein which can be in an active or inactive state (Fig. 3A). The protein is involved in two feedback loops: it promotes its own activation and inhibits its inactivation. The equation used to model this is

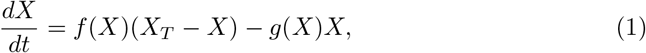

where *X* is the concentration of active protein and *X*_*T*_ is the total amount of this protein, *X*_*T*_ = *X* + *X*_inactive_. The functions *f* and *g* are given by

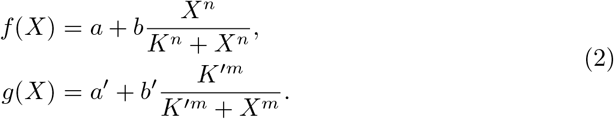

**Fig 3.**
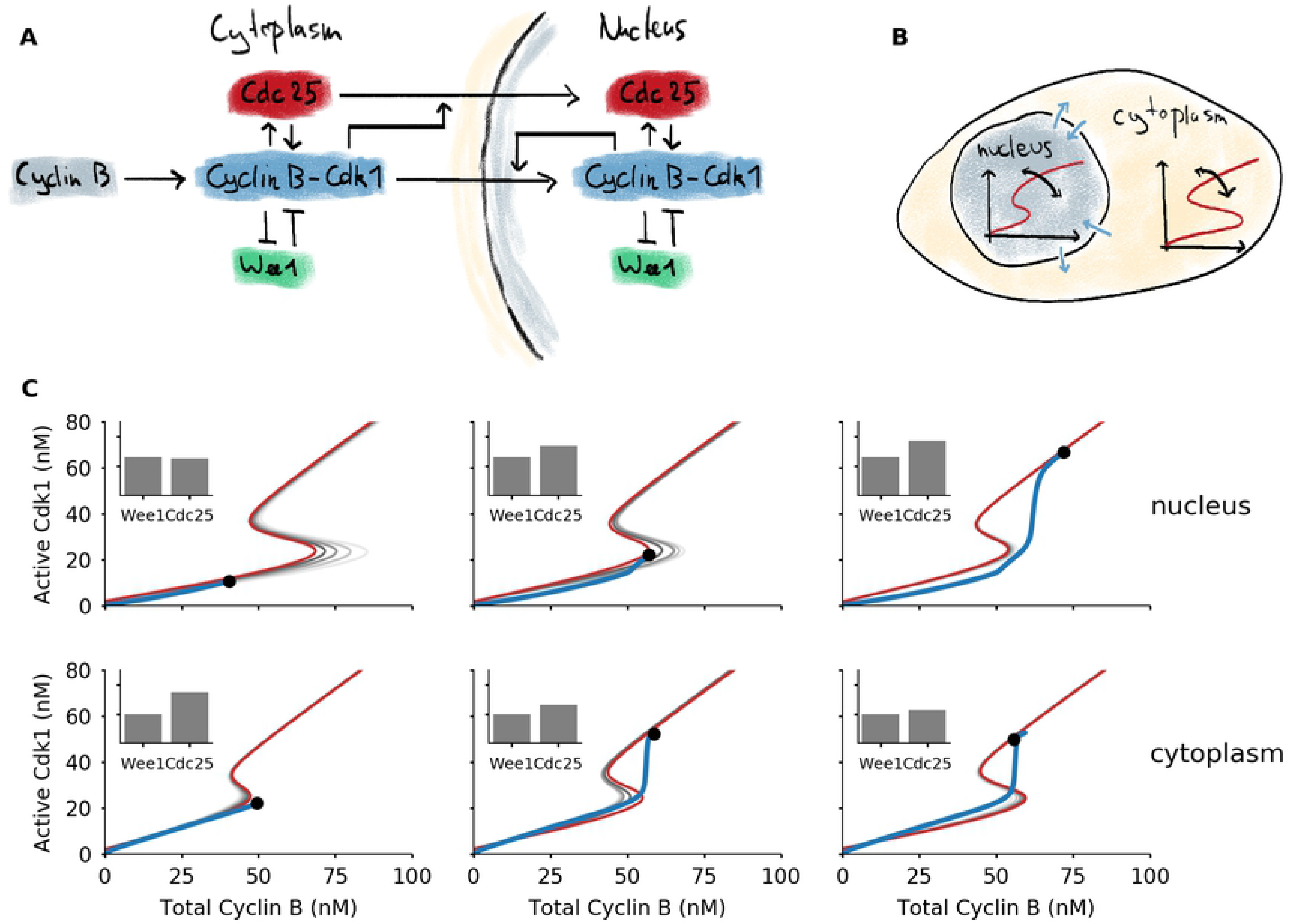
A simple model of protein activity shows bistability. A) Interaction diagram of a model, consisting of single protein which can be active or inactive. Active protein promotes its own activation and inhibits its inactivation. The basal activation rate is given by the parameter *a*. B) Steady state response to total protein *X*_*T*_, for different values of *a*. C) Steady state response to basal activation rate *a*, for different values of total concentration *X*_*T*_. D) Two-parameter bifurcation diagram of the system. The bistable region is shaded. The vertical orange curve, when followed from bottom to top, corresponds to the orange response curve Panel B. The horizontal green curve, followed from left to right, corresponds to the green response curve in Panel C. changes of the second bistable switch in mitosis. The effect of Greatwall and NEBD on the bistable response curves has been studied already in the context of mitotic collapse [57].

These response curves involve Hill functions, which are typically the outcome of basic biochemical reactions that generate ultrasensitivity, such as substrate competition, multisite phosphorylation, or others [58].

The combination of the different feedback loops and the steep response functions is known to generate bistability [11]. Indeed, this system shows bistable behavior, which can be visualized in different ways (Fig. 3B,C). The steady state response of the active protein level *X* can be bistable as function of the total amount of protein *X*_*T*_ (Fig. 3B). The shape of this response curve depends on the value of *a*, the basal activation rate of the protein. High values of *a* correspond to high basal activation of *X*. This ensures that any protein in the system will be directly converted into its active form, and *X* increases nearly linearly with *X*_*T*_. For low values of *a*, there is bistability, and the activation threshold becomes higher with lowering *a*. As a consequence, for very low *a* a large amount of protein needs to be added to the system to initiate the feedback loops that will lead to a full activation of the protein. If *X*_*T*_ is continuously increased, for example through a constant production of protein in the inactive state, at a certain moment the threshold will be reached and the system will jump to the active state. This representation is closely related to the cell cycle control system for mitotic entry, where the total abundance of Cyclin B (~ *X*_*T*_) gradually increases until the threshold for mitotic entry is reached and Cdk1 gets activated [23, 24, 33]. In this scenario, the activity of the phosphatase Cdc25 plays the role of the parameter *a* [35, 48].

A different bistable response emerges when plotting *X* as a function of *a*, keeping *X*_*T*_ fixed (Fig. 3C). The shape of the response curve now depends on the value of *X*_*T*_. For low values of *X*_*T*_, there is no bistability and the response is approximately hyperbolic. Intermediate levels of *X*_*T*_ lead to a bistable response curve. For high levels of *X*_*T*_, the switch becomes irreversible. In the latter case, the left threshold occurs at *a* < 0, which makes it impossible for the system to go back to its inactive state, since in biological systems *a* corresponds to an activation rate, a positive quantity. In such an irreversible switch, the system can transition from the low to high state, but it cannot go back. Dynamically varying *X*_*T*_ would provide a solution: by controlling the levels of *X*_*T*_, the transition back to the low state can be made possible. We have previously explored this mechanism in a model of the interaction between Cdk1 and the protein kinase Aurora B, which plays an important role during chromosome segregation in mitosis [59].

The effect of *a* and *X*_*T*_ can be summarized in a two-parameter bifurcation diagram (Fig. 3D). A response curve where only one of the two parameters is varied (Fig. 3B,C) corresponds to a horizontal or vertical cut in this diagram. In the remainder of this work, we will focus on the situation as in Fig. 3B: *X*_*T*_ is the main parameter – the input, as we previously called it – and we will explore the effects of having either a constant value of *a* or a dynamically changing *a*.

### 2.3 Transition timing is more robust and accurate in a system with a dynamic switch

In order to study the transition from low to high activity when protein is produced, we study the following system of equations:

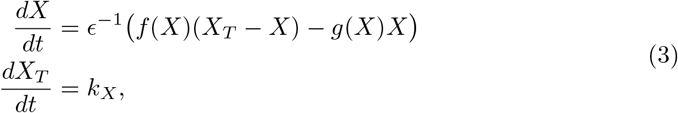

where the second equation corresponds to a constant increase in protein abundance *X*_*T*_. The small parameter *E* is added to model time-scale separation: the activation-inactivation dynamics of the protein are much faster than its production. As the total concentration increases, the system moves along the bottom branch of the bistable response curve. When the concentration crosses the activation threshold, the protein is rapidly activated (Fig. 4A).

**Fig 4.**
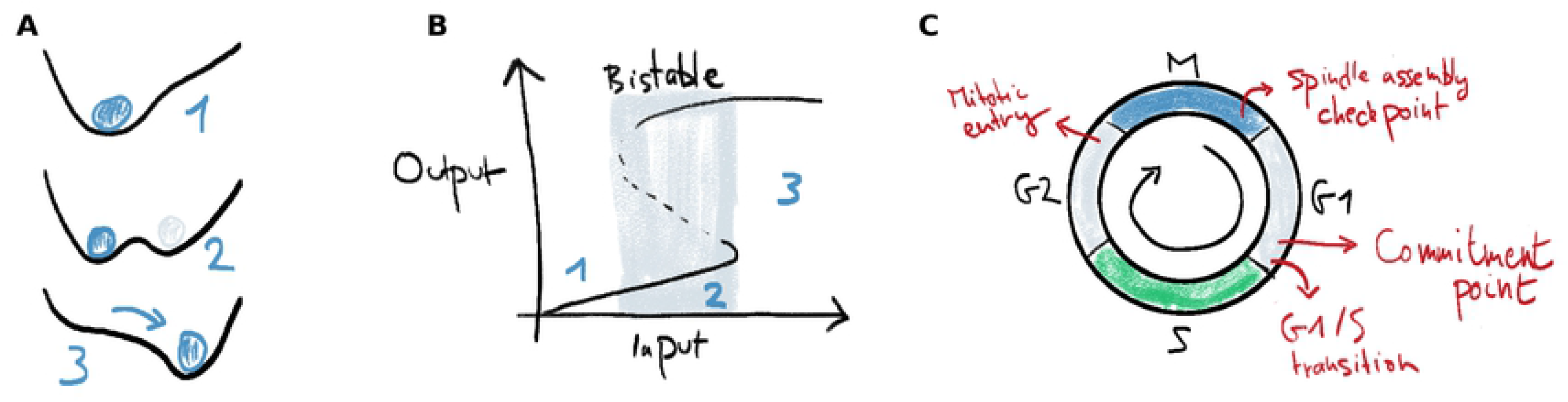
A dynamically changing bistable switch enhances robustness and accuracy of transitions in time and space. A) When *X*_*T*_ increases at a constant rate (here *k*_*X*_ = 0.2), the activity of the protein will increase suddenly at the moment *X*_*T*_ crosses the activation threshold of the switch. B) Function used to make the shape of the switch dependent on the total amount of protein, by coupling *a* to *X*_*T*_. The switch variation depends on the value of Δ*a*, which controls the magnitude of possible deviations of *a* from a mean value 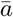. The parameter *κ* controls how abruptly the system switches between low and high *a* values. C) Time evolution of a system in which *X*_*T*_ increases at a constant rate, and *a* is coupled to *X*_*T*_. The gray response curves are snapshots in time. The activation threshold starts out to the far right, and moves left as *X*_*T*_, and with it *a*, increases. Here 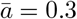, Δ*a* = 0.2, *κ* = 5, *k*_*X*_ = 0.2. D) Evolution of a system in which noise is added to the *X* variable. The transition time *t*_*c*_ is defined as the time when *X* crosses a threshold value. Here 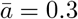, Δ*a* = 0.2, *κ* = 5, *k*_*X*_ = 0.2, *σ* = 0.6. E) Histogram of measured transition times for a static switch (Δ*a* = 0) and a dynamic switch (Δ*a* = 0.3), with 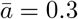, *κ* = 5, *k*_*X*_ = 1, *σ* = 0.6. The spread is lower for the dynamic switch. F) Coefficient of variation (CV), defined as standard deviation divided by mean, of the transition time, as function of the switch variation Δ*a*. Here 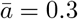, *κ* = 5. G) Coefficient of variation as function of *κ*, which defines the speed by which *a* changes. Faster changing corresponds to smaller deviations. Here 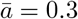, Δ*a* = 0.3. H) Velocity of a bistable front as function of *a*, for *X*_*T*_ = 2. A positive velocity means that the active protein state overtakes the inactive state (the front shown in panel I moves to the right). At *a* ≈ 0.15, the front is stationary. I) A bistable front in the presence of an inhomogeneous *a* profile in space. On the left, *a* = 0.27, which means the front moves to the right. On the far right, *a* = 0.13 is low, which means that the front moves to the left. The result is that the front is pinned in the middle where *a* ≈ 0.15. This pinning can be lifted by a redistribution of *a* (see S2 Video). Other parameters used in all simulations: *a*′ = 0.1, *b* = *b*′ = 1, *K* = *K*′ = 1, *m* = *n* = 5, *ϵ* = 0.05. Histograms and coefficients of variations where calculated over 200 simulations.

We then set out to investigate how this activation is affected when the shape of the bistable switch is changing while *X*_*T*_ is increasing. We impose the following functional form on *a* (Fig. 4B):

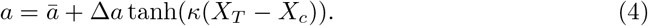

The value of 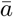 is the value around which *a* varies symmetrically. By tuning the parameter Δ*a* we can control the extent of the shape changes: for Δ*a* = 0, the switch does not change and 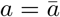 is a constant. For 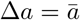, *a* varies between extremes of 0 and 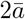. The parameter *κ* controls the abruptness with which the bistable shape changes when *X*_*T*_ crosses a threshold *X*_*c*_. This threshold is chosen to be in the middle of the bistable switch. For positive values of Δ*a*, the activation threshold of the switch moves to the left while *X*_*T*_ increases. The dynamics of such a system are illustrated in Fig. 4C and in S1 Video.

One striking consequence of the dynamic switch is that the level of *X* is kept very low until its activation, whereas if *a* is not changing, *X* already increases while the system is approaching the threshold. Moreover, by lowering the activation threshold while the system is approaching the transition point, the timing of activation can be controlled more precisely. To illustrate this, we add noise with magnitude *σ* to the *X* variable (see Methods for details). As a result, noise can trigger the system to jump to the high activity state even before the activation threshold is reached. We simulate the system many times and measure the transition time, defined as the time the value of *X* crosses a threshold (Fig. 4D). A dynamic switch shows less variation in the transition time than a static switch (Fig. 4E). Increasing the amplitude (higher Δ*a*) and abruptness (higher *κ*) of the dynamical shape changes help to further decrease the variation in transition timing (Fig. 4F-G).

Noise is inevitable in biochemical systems, and can be a nuisance or something the cell uses to its advantage [60]. Here, in the context of the cell cycle, premature activation due to noise is to be avoided. We conclude that accurate control of the timing of transition can be achieved by dynamically changing the switch and increasing *a* as *X*_*T*_ approaches the threshold. Note that a more realistic description of stochasticity would require using a master-equation or Fokker-Planck approach. However, our current approach, using a Langevin equation, is sufficient to demonstrate the utility of a dynamically changing switch (see Methods section).

### 2.4 Transitions in space can be controlled by dynamically changing the bistable switch

When bistable systems are coupled in space in the presence of diffusion, they may produce traveling fronts. In our model with active and inactive protein, a traveling front can arise when one region of space has a high *X* activity and an adjacent region has low activity. The interface between these regions starts to move, depending on which state is dominant. Such traveling fronts are omnipresent in biology, where they usually have a signalling or synchronizing function [42, 61].

The speed and direction of traveling fronts depend on the parameters of the system. Consider for example a traveling front that links regions of high and low activity of the protein *X*, with *X*_*T*_ = 2 fixed. The direction of the front depends on *a*: low *a* corresponds to a dominant low activity state, high *a* to a dominant high activity state. The front moves such that the dominant state overtakes the other one (Fig. 4H).

Let us assume that the parameters may vary in space, such that the front speed itself varies in space. Consider a system where *a* is high in one region, low in another and has a smooth transition between both regions. In this case, the front would move to the right until it hits the transition region, where it slows down and comes to a halt (Fig. 4I). This phenomenon is called pinning. Front pinning and localization – possibly due to a spatial inhomogeneity, as here – occur frequently in physical systems [62]. In biology, front pinning mechanisms have been studied in the context of cell polarization [63], and in ecosystem transitions [64].

The front comes to a halt due to a spatially heterogeneous profile of *a*. A pinned front can then be released by redistributing *a*, and thus changing the bistable switch and the dominant state (see S2 Video). Dynamically changing the parameters which affect the shape of the bistable response curve can thus provide the cell with extra control over spatial transitions.

Waves of Cdk1 activity spread throughout the cell at mitotic entry, as has been observed in *Xenopus* cell-free extracts [65, 66] and in the early *Drosophila* embryo [67, 68]. In the cell, and in extracts, spatial heterogeneities are present through nuclei, which concentrate certain proteins [66]. In such systems, the effect of dynamic bistability on front dynamics is likely present, all the more because the spatial heterogeneity drastically changes at nuclear envelope breakdown. In *Drosophila*, dynamic changes in the bistable switch have been shown to play an important role in determining the nature of mitotic waves [68].

### 2.5 A dynamic switch promotes stable oscillations

Fast transitions between states form the basis of relaxation oscillations, such as those observed in early embryonic cell cycles of *Xenopus laevis*, where the cell quickly switches between interphase and mitosis. We asked ourselves how such oscillations are affected by a dynamic bistable switch. We expanded our model to obtain oscillations in protein activity and abundance by including production and degradation:

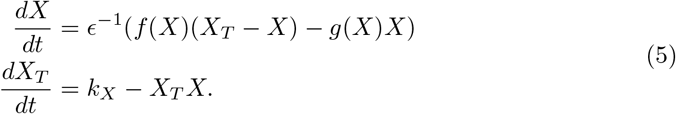

As before, *k*_*X*_ is the protein production rate and the dynamics of activation and inactivation are fast with respect to production and degradation, such that Ε is small. We assume that the active form of *X* promotes its own degradation through mass-action kinetics (Fig. 5A). Note that we have simplified the set of equations by omitting a term (−*X*^2^) from the first equation. In doing so, we ensure that the bistable response curve we have used before appears as a nullcline of the system. This simplification does not significantly change the system dynamics (see the Methods section).

**Fig 5.**
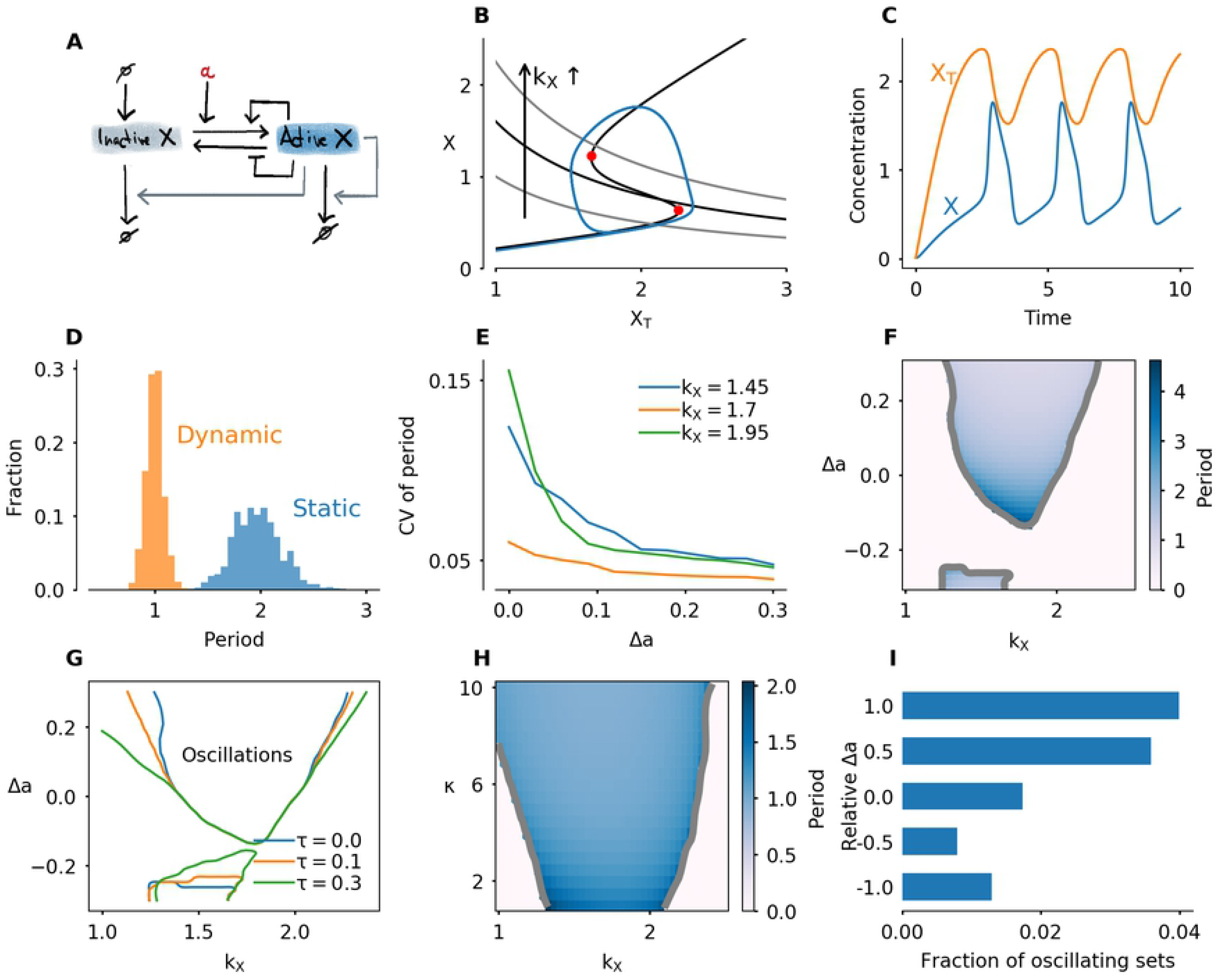
Dynamic switches promote oscillations. A) Interaction diagram. The active form of protein *X* promotes its own degradation. B) Phaseplane of the system given by Eq. 5 (static switch). The second nullcline depends on the value of *k*_*X*_. The S-shaped nullcline has the same shape as the bistable response curve studied in the previous section. The second nullcline is given by *X* = *k*_*X*_/*X*_*T*_ and is shown for *k*_*x*_ = 1, 1.6 and 2.25. The blue limit cycle corresponds to *k*_*X*_ = 1.6. C) Time series of the oscillatory system with *k*_*X*_ = 1.6. D) Histogram of the period for a static (Δ*a* = 0) and dynamic (Δ*a* = 0.3) switch, with 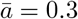, *κ* = 5, *σ* = 0.4. Simulation for a total time of *T* = 2000. E) Coefficient of variation (standard deviation divided by mean) of the period in the oscillatory system with noise added to the *X*-variable. Here *κ* = 5, *σ* = 0.2. F) Period in color, as function of *k*_*X*_ and Δ*a* with *κ* = 5, *τ* = 0, 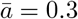. G) Oscillatory region in the (*k*_*X*_, Δ*a*)-plane for different values of the delay time *τ* with *κ* = 5, 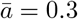. H) Period as function of *k*_*X*_ and *κ* with 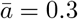 = 0.3, Δ*a* = 0.3 and *τ* = 0.1. I) Fraction of parameter sets for which the system oscillates for 10000 randomly sampled parameter sets. Other parameters used in all simulations except for Panel I: *a*′ = 0.1, *b* = *b*′ = 1, *K* = *K*′ = 1, *m* = *n* = 5, *ϵ* = 0.05.

For given functions *f* and *g*, this system only oscillates for a specific range of *k*_*X*_. If *k*_*X*_ is too small, the production rate is not high enough to push the system over the activation threshold, and the system converges to a steady state with low activity. For a high value of *k*_*X*_, degradation cannot compensate for production even when the protein is mostly active, and the system converges to a steady state with large *X*. For intermediate values of *k*_*X*_, the system switches between accumulation and degradation with low and high *X* respectively. This is illustrated in the phase plane in Fig. 5B. If *k*_*X*_ is such that the two nullclines intersect in between the two saddle-node points, the system converges to a stable limit cycle. This oscillation is marked by a slow increase along the bottom branch of the bistable curve, a slow decrease along the upper branch, and fast jumps in between (Fig. 5B,C).

As before, we allow the parameter *a* to depend on the total amount of protein (Eq. (4)). In the previous section we found that, in the presence of noise, transition times show less variation if the switch dynamically changes. In the case of oscillations, we find that more accurate transition times are reflected in a more stable period. A dynamic switch (Δ*a* > 0) shows less variation in the period of the oscillation than a static switch (Fig. 5D). Larger switch changes ensure smaller variation of the period (Fig. 5E), and this effect is more pronounced for extreme values of *k*_*X*_, such that the nullclines intersect close to the saddle-node point. For those values, stochastic activation/inactivation is more likely, an effect which is mitigated by the dynamically changing switch. In *Xenopus laevis*, early embryonic cycles have a remarkably stable period [69]. Combined with an initial period difference between different cells in the embryo, this shows as a wave of cell division. It is possible that dynamically changing bistable switches contributes to such stability in periods.

Next, we expanded the model by also looking at negative values of Δ*a*, which corresponds to an activation threshold which increases as *X*_*T*_ increases. Moreover, we model a biologically plausible phase shift to the relation between *X*_*T*_ and *a* by including a time delay *τ* :

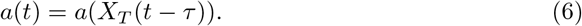

As mentioned before, oscillations occur if *k*_*X*_ lies in a given interval. This interval becomes larger if Δ*a* increases, which indicates that larger changes of the bistable curve produce a larger region of oscillations (Fig. 5F). This observation holds both with and without time delay, but the effect is larger if time delay is included (Fig. 5G). The effect of increasing time delay is most pronounced for low *k*_*X*_. Note that also for Δ*a* < 0, there is an oscillatory region. When Δ*a* < 0, the switch changes in the opposite direction of the change of *X*_*T*_, i.e. the activation threshold moves to the right as *X*_*T*_ approaches it. The speed by which the bistable response curve changes also plays a role: faster, more abrupt transitions, which correspond to higher *κ*, promote oscillations (Fig. 5H).

As a final demonstration of how dynamically changing switches affect the occurence of oscillations, we perform a random sampling of 10000 parameter sets. We sample all the parameters affecting the Hill functions *f* and *g*, the timescale parameter *ϵ*, and *k*_*X*_. For each parameter set we first detected whether the system is bistable, and if so, we simulated the model with Δ*a* = *ia, i* = −1, −1*/*2, 0, 1*/*2, 1, for *κ* = 1, 5, 10 and *τ* = 0, 0.1, 0.2. For each simulation we detect whether the system oscillates or goes into a steady state. Generally, oscillations are quite rare, but in all cases, having Δ*a* > 0 increased the probability of obtaining oscillations (Fig. 5I and Figure in S1 Supporting Information). Note that oscillations can also exist for Δ*a* < 0. In that case, however, oscillations are less likely.

To conclude, we have found that making the bistable response curve dynamic instead of static enhances the accuracy of the oscillation period in noisy systems, and increases the region in parameter space where oscillations are found. This effect is larger when the shape change lags the increase of *X*_*T*_. S3 Video shows an oscillation with a dynamically changing bistable switch.

## 3 Discussion

Bistable switches play a crucial role in the cell cycle, providing a mechanism for quick and irreversible transitions. They also lie at the basis of more complex behavior such as spatial front propagation and relaxation oscillations. The classic viewpoint of a static switch and fixed activation and inactivation thresholds does not take into account that the factors that determine the shape of the response curve can vary over time. In a biochemical system, these factors are typically protein concentrations. When those concentrations change, the bistable response curve changes, and this happens all while the system is proceeding along the branches of the bistable curve.

By making this dynamically changing bistability explicit in a simple model, we have shown that such a mechanism allows more accurate control of the transition timing in noisy systems. Additionally, by controlling protein levels in space, the location and speed of propagating fronts of activity can be regulated. In oscillatory systems, a changing bistable switch increases the robustness of the oscillations to parameter variations. The control of transition timing and avoiding a premature transition can play a role in cell cycle checkpoints, whereas the enhanced oscillations may be important in embryonic cell cycles which behave like autonomous oscillators. These advantages suggest that such dynamic regulation may have evolved as an extra mechanism in the cell’s repertoire to ensure robust faithful genome replication and division.

Besides accuracy and robustness, a dynamic bistable switch may provide other benefits to the cell. If there is an energetic cost associated to maintaining a bistable switch at a certain level, dynamically controling the shape can be a way to more efficiently use energy. This kind of temporal compartmentalization is widely seen in biology. Circadian rhythms, for example, provide a means of compartmentalizing processes to align with external light and temperature cycles and therefore optimally use energy [70]. This energy-based view of dynamic bistable switches will perhaps benefit from a thermodynamic description, which takes into account energy consumption (e.g. [71]).

We have shown that not only temporal variation of the response curve, but also spatial control can play a role. We demonstrated how the location of a traveling front can be controlled by modifying the bistable switch in space and time. The biological example of mitotic entry shows that compartmentalization in space can add dynamics which are not seen in a well mixed system. Having two compartments with slow fluxes between them essentially splits the system in two: two bistable response curves now determine the evolution of the system, and their shape can be tuned by shuttling proteins around. This can be used to obtain different activation thresholds in different compartments and thereby add extra control over mitotic entry.

Spatial compartmentalization of biochemical reactions has recently become a topic of interest for mathematical modelers, since it can add new dynamics to otherwise well-mixed systems. A monostable system may become bistable by adding compartments [37]. This increases the complexity and richness of the system under study: by compartmentalizing chemical reactions, cells can locally increase concentrations to speed up reactions, or inversely keep certain reactions from happening by separating the reactants. Adding multiple compartments is also attempted in synthetic biological systems by introducing artificial membranes [72, 73].

In experiments too, compartmentalization and spatial organization have been found to introduce new dynamics. Santos et al. [36] showed that spatial feedbacks are present in mitotic entry, which we used as one of the mechanisms in our biological example. Recently Doncic et al. showed that compartmentalization of a bistable switch plays a role in the commitment point, also referred to as the *Start* checkpoint in yeast [74].

We believe that mathematical modeling of these spatial aspects of cell cycle transitions is a fruitful way to extend our insights into cell cycle regulation. Numerous mathematical models of the cell cycle already exist. Spatial regulation, however, is mostly absent. Models that include a spatial component often focus on traveling waves which can play a role in synchronizing large cells, such as *Xenopus* embryos [65] or *Drosophila* syncytia [67]. In *Xenopus* cell-free extracts, nuclei play an essential role as a pacemaker, possibly due to the fact that nuclei locally increase concentrations of key regulatory proteins, in turn changing the bistable switch upon which mitotic entry is built [66]. In *Drosophila*, changes to the bistable switch have been proposed as an explanation of changing wavespeeds over different cycles [67], and bistable thresholds play a crucial role in the so-called sweep waves [68]. New insights are likely to be gained from models that also take into account the heterogeneity and spatial structures of a real cell. A first step towards that goal is to consider the compartments of nucleus and cytoplasm. Some models have done this already – e.g. in the context of cell growth [75] – but we believe there is a vast range of new biological insights to be gained by studying new kinds of mathematical models in that direction. The viewpoint of changing bistable switches may play a helpful role in analyzing such models.

From the mathematical standpoint, figuring out how such spatial heterogeneity is best introduced poses an interesting challenge, where simplicity, computational efficiency and realism have to be weighed against one another. The method we used in the model of mitotic entry was ODE based. For each compartment, every chemical species has its own ODE. This assumes that, inside a compartment, diffusion is fast and the system is well mixed. Another option is to use fully spatial models consisting of partial differential equations (PDEs). Here, boundary conditions should be used to model fluxes between different compartments. Yet another type of model is hybrid: some small compartments are considered to be well-mixed, and are modeled by ODEs, whereas transport through the medium between the compartments is governed by a PDE (see e.g. [76] for a recent example of such a model in a biological system). If compartments are not bound by membranes, but instead generated by phase separation, modeling may need to take into account the physics of the phase separation process, a topic of current interest in cell biology [77].

The implementation of the changing response curve we used here – with the sigmoidal function *a*(*X*_*T*_) and the time lag *τ* – is artificial but suitable for the message we want to convey. However, in the future a more thorough theoretical study of such simple models could replace the explicit dependence of *a* on *X*_*T*_ by evolution equations for the dynamics of the switch shape. It would be valuable to study a model of the form

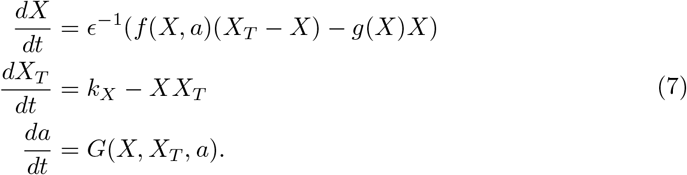

This equation takes the form of a two-slow, one fast dynamical system: the activation dynamics of *X* happen on a faster scale than the production and degradation of *X*_*T*_ and the dynamics of *a*. Rigorous mathematical study of such multiple-timescale systems has been done extensively in other areas, such as mathematical neuroscience [78, 79]. Connecting a more rigorous mathematical analysis to expected biological outcomes in the cell cycle can provide important clues to uncover the underlying dynamics, and may lead to new opportunities to connect mathematicians and cell biologists.

Experimental observations of dynamically changing bistable switches can take different forms. One of the outcomes of a mathematical model such as the one we studied here is a time series, which gives the evolution of concentrations of the main proteins over time. By performing a more detailed analysis, we can find out which qualitative features are specific to a timeseries derived from changing bistability. Next, experiments can be set up to try to detect such features. Another approach would be to measure the steady-state response curves to obtain activation thresholds, and perform this experiment under different experimental conditions.

Mathematical modeling and concepts derived from nonlinear dynamics, such as bistability and limit cycles, have been very influential on our understanding of many biological phenomena, and will continue to be, as has been recently advocated by Tyson and Novák [80]. In our discussion we have echoed some of their perspectives.

Additionally, in this paper, by adding an extra layer to the regulation of cell cycle transitions we have attempted to push our dynamical understanding of this fundamental process a little bit further.

Even though the simple model we studied in this paper was artificial, its main conclusions will likely hold for more realistic mathematical models. In fact, we suspect that a dynamically changing bistable switch is already present in many published models, but not recognized or described as such. Therefore, we propose that the dynamically changing switches we described can be used as a means to interpret existing models and perhaps inspire new ones.

## 4 Materials and methods

### 4.1 Sofware and algorithms

Simulations, data analysis, plotting and animations were all done in Python except for the two-parameter bifurcation diagram in Fig. 3, which was created using the interface to AUTO of the software XPPAUT [81]. The majority of our simulations use ODEs, but some versions of the model are delay differential equations or stochastic differential equations. To simulate all of these, we made use of the software packages JITCODE, JITCDDE and JITCSDE for Python, which implement solvers for ordinary, delay and stochastic differential equations respectively [82]. For parameter sweeps we used a high-perfomance computing cluster.

To obtain the bistable response curve, we implemented a pseudo-arclength continuation algorithm directly in Python (see, e.g., [83]). This gave us the flexibility to compute response curves on the fly, as for the animations.

For the period detection in the oscillatory systems, we start from the time series *X*(*t*) and detect the times *t*_*u,i*_ and *t*_*d,i*_ when *X* crosses a certain threshold up and a certain threshold down. The threshold up is the vertical coordinate of the leftmost saddle-node point, the down threshold is the vertical coordinate of the rightmost saddle-node point. Next, we compute the period as the difference *t*_*u,i*+1_ − *t*_*u,i*_. In the noisy system, this gives a set of period *P*_*i*_ on which we can perform statistics. For the deterministic systems this value is constant. This method ensures that we only track oscillations that go around both branches of the bistable system, i.e. of sufficient amplitude.

To detect the contours of the oscillatory regions in Fig. 5F-H, we detected all points in the heatmap where the period goes from zero to positive and then applied smoothing. Note that this boundary is not strictly the same as the boundary between steady state and oscillations, since we consider only oscillations of sufficient amplitude, that go around both branches of the switch.

Our code and demo files that show how to use it are available at https://github.com/JanRombouts/dynamicswitches.

### 4.2 Stochastic model

The stochastic equations we use are

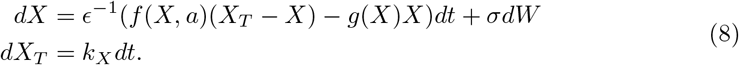

These equations are of Langevin type with noise only in the fast variable, which is not a correct representation of molecular noise, but the simplest way to extend our ODE model to include stochasticity. We include noise only in the fast variable to simplify the system and only allow transition through ‘vertical’ deviations from the steady state branch, in line with typical studies on stochastic switching [84]. Moreover, the ratio of noise magnitude in fast and slow variable is high when stochastic differential equations are derived from a discrete stochastic model [85], such that setting the noise on the slow variable to zero is reasonable. To determine transition times, we detect the timepoint when *X* crosses a threshold concentration. This threshold concentration is always the average vertical coordinate of the saddle-node points of the static bistable switch.

### 4.3 Removal of degradation term in the X-equation

In the oscillatory system, *X* degrades itself, both in active and inactive form. The set of equations corresponding to this is

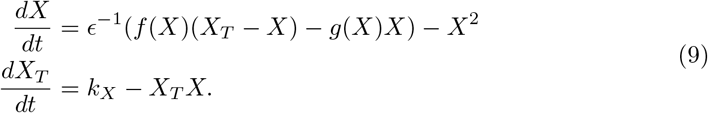

We can rewrite this as

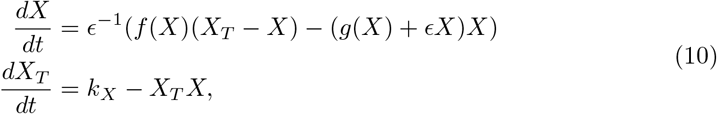

and since *ϵ* is considered to be small, the shape of the bistable switch induced by these equations is nearly the same as that induced by the one where *ϵ* = 0, which we use.

### 4.4 Spatial model

For the simulations in space, we use the equations

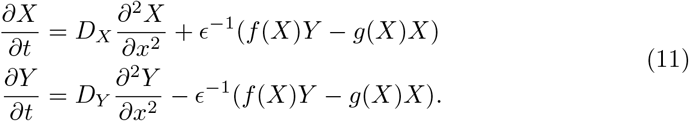

here, *Y* is the inactive form of the protein. These equations allow more flexibility in choosing, for example, different diffusion constants for active and inactive form. We took *D*_*X*_ = *D*_*Y*_ = 5 in our simulations for Fig. 4H,I. We simulated these equations using a forward difference in time and centered difference for the space derivative. We use zero-flux boundary conditions. For Fig. 4H, we detect the front position as function of time and fit a linear function. Initial conditions are always a step function.

### 4.5 Sampling of parameters

Sampling of the parameter sets in Fig. 5I was done as follows: each parameter was sampled uniformly and independently from a given interval. For *ϵ* we sampled the logarithm. In S1 Supporting Information we provide a table with the intervals and whether the parameter was sampled logarithmically or not. The system was simulated with sampled parameters for a total time of *T* = 200. We considered a set of parameters as oscillatory if the period is larger than 0.01.

### 4.6 Full set of equations for the biological example

We keep track of three different variables: total Cyclin B-Cdk1 complexes ([Cyc], active Cyclin B-Cdk1 complexes ([Cdk1]) and Cdc25 levels ([Cdc25]). The model equations are based on the equations used by Yang and Ferrell [56]. Note that Cdc25 is a scaled variable: the value 1 would correspond to the level assumed by Yang and Ferrell. Each variable has a nuclear and cytoplasmic version which are denoted by subscript *n* or *c*. Cyclin B is constantly produced at a rate *k*_*s*_ and binds immediately to Cdk1 to create the complex. We assume that production only happens in the cytoplasm. The activation rate of Cdk1 depends on Cdc25 levels and activity. The level is controlled by the variable [Cdc25], the activity is a function of Cdk1, since Cdk1 is an activator of Cdc25. The inactivation rate of Cdk1 depends on Wee1 levels and activity. We assume that total Wee1 levels are constant, but this level is higher in the nucleus than in the cytoplasm. In the simulation used in Fig. 2, we used [Wee1]_*n*_ = 1.3 and [Wee1]_*c*_ = 1. This variable is also scaled, [Wee1] = 1 corresponding to the model used by Yang and Ferrell. As initial conditions for Cdc25, we use [Cdc25]_*n*_ = 1, [Cdc25]_*c*_ = 2.

Cyclin B-Cdk1 complexes and Cdc25 can be imported and exported from the nucleus with certain import and export rates. We use the convention that the subscript *n* or *c* for the rate denotes the compartment towards which the protein is moved. To account for the observation that both Cyclin B-Cdk1 and Cdc25 import is increased at mitotic entry, we introduce the functions *I*_Cyc_ and *I*_Cdc_, which modify the import rates of Cyclin B-Cdk1 and Cdc25 respectively. We use

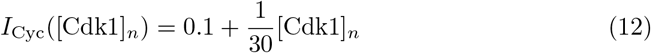

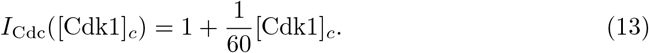

The equations are

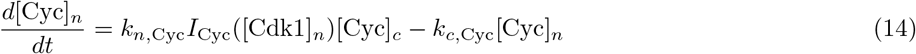

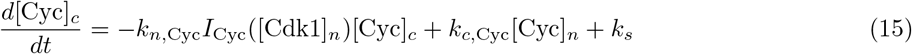

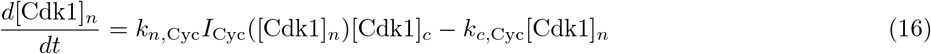

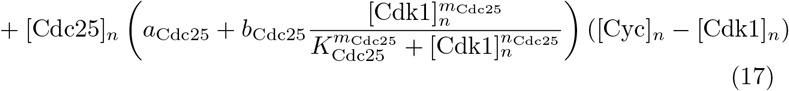

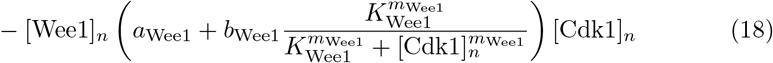

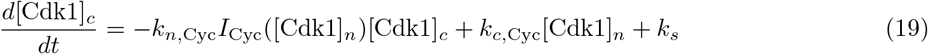

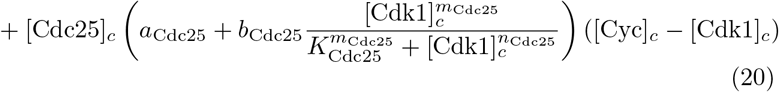

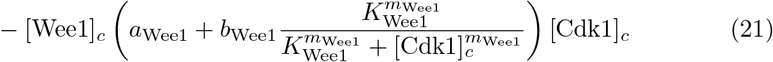

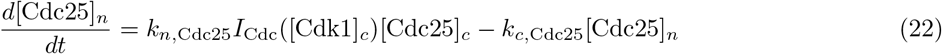

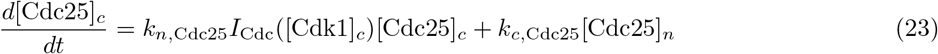

The parameters can be found in S1 Supporting Information, and are mostly taken from [56]. The import rates and the functions that influence those rates were chosen to obtain a good example of the mechanism we propose.

## Supporting information

**S1 Supporting Information. File containing two tables and one extra Figure.** Table with parameter values used in the cell cycle model, table with the parameter bounds used in the sampling, figure with the fractions of oscillatory parameter sets for different values of *τ* and *κ*.

**S1 Video. Transition with a changing switch.** This animation corresponds to Fig. 4C. The protein is produced at a constant rate, while the activation threshold is moving to the left due to an increase in *a*. The effect is a fast transition, and *X* activity stays low until the transition.

**S2 Video. Moving front with redistribution of *a*.**

This animation illustrates the blocking of the front due to a heterogeneity in *a*, and the release of the front due to redistribution of *a* (Fig. 4H,I). The *a* profile we use is a smooth hyperbolic tangent function of *x*. The high value of *a* is 0.27, the low value is The front gets stuck at the transition to low *a*, where *a* ≈ 0.15. At time *t* = 25, we effect a smooth transition to the flipped profile which releases the front, after which it continues moving to the right.

**S3 Video. Oscillation with a changing switch.**

This animation illustrates a system with production and degradation. We took *k*_*X*_ = 1., 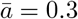, Δ*a* = 0.2, *κ* = 5, *τ* = 0. The bistable switch is changing while the system oscillates.

**S4 Video. Mitotic entry with two compartments**

This animation corresponds to Fig. 2 in the main text. At first, Cyclin B accumulates in the cytoplasm. The activation threshold for Cdk1 is lower there, so Cdk1 activity jumps to the upper branch first in the cytoplasm. this triggers nuclear import of Cdc25, which lowers the threshold in the nucleus. Following this, Cdk1 activity in the nucleus jumps up, which triggers an increased import of Cyclin B. Cdk1 activity in the nucleus keeps increasing while in the cytoplasm it settles to a constant value.

## Acknowledgments

We are grateful to the members of the Gelens lab for useful comments on the manuscript. The computational resources and services used in this work were provided by the VSC (Flemish Supercomputer Center), funded by the Research Foundation - Flanders (FWO) and the Flemish Government – department EWI.

